# Acetyl-CoA carboxylase Inhibition increases RPE cell fatty acid oxidation and limits apolipoprotein efflux

**DOI:** 10.1101/2023.11.07.566117

**Authors:** Daniel T. Hass, Kriti Pandey, Abbi Engel, Noah Horton, Brian M. Robbings, Rayne Lim, Martin Sadilek, Qitao Zhang, Gillian A. Autterson, Jason M.L. Miller, Jennifer R. Chao, James B. Hurley

## Abstract

**Purpose:** In age-related macular degeneration (AMD) and Sorsby’s fundus dystrophy (SFD), lipid-rich deposits known as drusen accumulate under the retinal pigment epithelium (RPE). Drusen may contribute to photoreceptor and RPE degeneration in AMD and SFD. We hypothesize that stimulating β-oxidation in RPE will reduce drusen accumulation. Inhibitors of acetyl-CoA carboxylase (ACC) stimulate β-oxidation and diminish lipid accumulation in fatty liver disease. In this report we test the hypothesis that an ACC inhibitor, Firsocostat, limits the accumulation of lipid deposits in cultured RPE cells.

**Methods:** We probed metabolism and cellular function in mouse RPE-choroid, human fetal-derived RPE cells, and induced pluripotent stem cell-derived RPE cells. We used ^13^C6-glucose and ^13^C16-palmitate to determine the effects of Firsocostat on glycolytic, Krebs cycle, and fatty acid metabolism. ^13^C labeling of metabolites in these pathways were analyzed using gas chromatography-linked mass spectrometry. We quantified ApoE and VEGF release using enzyme-linked immunosorbent assays. Immunostaining of sectioned RPE was used to visualize ApoE deposits. RPE function was assessed by measuring the trans-epithelial electrical resistance (TEER).

**Results:** ACC inhibition with Firsocostat increases fatty acid oxidation and remodels lipid composition, glycolytic metabolism, lipoprotein release, and enhances TEER. When human serum is used to induce sub-RPE lipoprotein accumulation, fewer lipoproteins accumulate with Firsocostat. In a culture model of Sorsby’s fundus dystrophy, Firsocostat also stimulates fatty acid oxidation, improves morphology, and increases TEER.

**Conclusions:** Firsocostat remodels intracellular metabolism and improves RPE resilience to serum-induced lipid deposition. This effect of ACC inhibition suggests that it could be an effective strategy for diminishing drusen accumulation in the eyes of patients with AMD.

## Introduction

Age-related macular degeneration (AMD) is a retinal degenerative disease and the leading cause of blindness in older adults (1). The ultimate cause of degeneration in AMD is unknown, but age, smoking, and circulating HDL-cholesterol are among the risk factors (2). Photoreceptor degeneration in AMD is associated with accumulation of soft drusen, basal laminar deposits, and subretinal drusenoid deposits (3). These deposits are rich in lipids (4–8) and lipoproteins (9–11), and are thought to arise from retinal pigment epithelium (RPE) cell components (12). The hydrophobic nature of these lipid-rich structures may impede transport of hydrophilic metabolic fuels like glucose through the RPE to the retina (13). Diminished nutrient transport may contribute to photoreceptor degeneration in AMD (13). Soresby fundus dystrophy has a similar pathology as AMD, but with an earlier onset of symptoms (14, 15).

If a hydrophobic barrier generated by drusenoid structures compromises photoreceptor viability, removing it could slow or stop degeneration (16–18). Our goal is to limit the formation of these lipid-rich structures by stimulating fatty acid degradation. A decrease in fatty acid levels should limit the amount of lipid and lipoprotein available to form drusen.

Fatty acid synthesis and oxidation are controlled by malonyl-CoA. Malonyl-CoA is a product of acetyl-CoA carboxylases (ACCs) (19). Two isoforms of ACC control fatty acid and lipid homeostasis. Malonyl-CoA produced by cytosolic ACC1 is a substrate for fatty acid synthesis (19, 20). Malonyl-CoA synthesized on the mitochondrial outer membrane by ACC2 prevents oxidation of long-chain fatty acids in mitochondria by inhibiting carnitine palmitoyl transferase (CPT) (21). Esterification of long chain fatty acids to carnitine by CPT is required for fatty acid import and oxidation in the mitochondrial matrix (22).

ACCs must dimerize to produce malonyl-CoA. Firsocostat (also referred to as GS-0976 or ND- 630) is a small molecule that inhibits dimerization of both ACC isoforms (23). It limits malonyl-CoA formation, decreases circulating lipid levels, and decreases hepatic lipid accumulation.

Firsocostat and other ACC inhibitors are being pursued as a potential therapeutic for non-alcoholic fatty liver disease (23–26). Our goal is to determine whether inhibition of ACCs limits production or accumulation of lipoproteins adjacent to RPE.

We investigated the impact of Firsocostat-mediated ACC inhibition on fatty acid oxidation, lipid levels, apolipoprotein transport and deposition in mouse RPE-choroid and human RPE cells. Firsocostat increases the rate of fatty acid oxidation, remodels lipid composition, and decreases apolipoprotein export by the RPE. Several different ACC inhibitors have been developed and are in clinical trials for the treatment of non-alcoholic fatty liver disease. Our data suggest that in addition to treating liver disease these inhibitors could potentially limit a pathological increase in drusen during AMD.

## Methods

### Mice

All experiments adhered to our approved IACUC protocol and the ARVO statement for the use of animals in ophthalmology research. For animal experiments, we used 2-6 month-old wild-type C57BL/6J mice (JAX, stock# 000664) fed ad libitum on a 12-12 light/dark cycle. Mice were euthanized by awake cervical dislocation. The time of day for euthanasia was later than 11 pm, to avoid variability due to peak outer segment phagocytosis. To obtain RPE-choroid tissue for experiments, eyes were first dissected from the head into room temperature Hank’s buffered salt solution (Gibco, 14170120). Eyes were cleared of extraocular muscle and collagenous tissue, then cut along the ora serrata. Corna, iris, and lens tissue were removed, then the remaining retina and RPE-choroid were teased apart for downstream experiments. This process typically takes ∼3-4 minutes for both eyes from one mouse.

### Measurement of fatty acid oxidation – mouse RPE-choroid

Freshly dissected retina and RPE-choroid were incubated in Krebs-Ringer-Bicarbonate buffer (KRB) containing 5 mM D-glucose, 50 µM [U-^13^C16]-palmitic acid (Cambridge isotope labs, CLM- 409) complexed to bovine serum albumin at a 6:1 ratio of fatty acid to albumin, and vehicle (ethanol) or Firsocostat (Caymen chemical, 23961). Incubation medium was equilibrated >30 minutes at 37°C and 5% CO2 prior to the tissue dissection. Tissues were incubated for 1 hour then flash-frozen in liquid N2 and stored at -80°C prior to downstream processing.

### Measurement of fatty acid oxidation – RPE cells

RPE cells were incubated in pH 7.4 α-MEM and 50 µM [U-^13^C16]-palmitic acid (Cambridge isotope labs, CLM-409) complexed to bovine serum albumin at a 6:1 ratio of fatty acid to albumin. This buffer was pre-equilibrated at 37°C, 21% O2, and 5% CO2 prior to incubations. We changed culture medium on cells, washed them in sterile 1x PBS, and incubated them in medium containing the ^13^C label. To determine metabolite uptake or export rate we sampled incubation medium at times indicated in the text and figures.

### Metabolite Extraction

Metabolites were extracted from tissue or media samples using 80% MeOH, 20% H2O, with 10 μM methylsuccinate (Sigma, M81209). The extraction buffer was equilibrated on dry ice, then 100-150 μL was added to each sample. Tissues were disrupted by sonication and incubated on dry ice for 45 minutes to precipitate protein. Proteins were pelleted at 17,000 x g for 30 minutes at 4°C. The supernatant containing metabolites was lyophilized at room-temperature until dry and stored at -80°C until derivatization.

### *Metabolite* Derivatization

Lyophilized samples were derivatized by adding 10 μL of 20 mg/mL methoxyamine HCl (Sigma, 226904) dissolved in pyridine (Sigma, 270970) and incubating at 37°C for 90 minutes. Samples were further derivatized by adding 10 μL tert-butyldimethylsilyl-N-methyltrifluoroacetamide (Sigma, 394882) and incubating at 70°C for 60 minutes.

### Gas Chromatography-Mass Spectrometry

Derivatized samples were injected into an Agilent 7890/5975C GC-MS system. Selected-ion monitoring was used to determine metabolite abundance, as previously described (27). Peaks were integrated using MSD ChemStation E.02.01.1177 (Agilent Technologies), and correction for natural isotope abundance was performed with IsoCor, v1.0 (28). Corrected metabolite signals were converted to molar amounts by comparing metabolite peak abundances in samples with those in a ‘standard mix’ containing known quantities of metabolites we routinely measure. Multiple concentrations of this mix were extracted, derivatized, and run alongside samples in each experiment. These known metabolite concentrations were used to generate a standard curve that allowed for metabolite quantification. Metabolite abundance was normalized to tissue protein concentration, and following this, paired tissues such as retinas and RPE- choroid from the same mouse were treated as technical replicates and averaged.

### Lipidomics - extraction

Mouse RPE-choroid was dissected then incubated in KRB supplemented with 50 µM unlabeled palmitate-BSA and 5 mM glucose, at 37°C and 5% CO2 for 6 hours. Tissue was flash-frozen in liquid nitrogen. To extract tissue lipids, samples in purified LC/MS-grade water were homogenized in a Bullet Blender with zirconia beads for 15 minutes then transferred to glass culture tubes. 25 μL of an isotope labeled internal standard mixture, 575 μL of methyl-*tert*-butyl ether (MTBE) and 160 μL of methanol (MeOH) were added to each sample. Samples were shaken for 30 min at room temperature. 200 μL of H2O was added, then samples were centrifuged at 2,500 x g for 3 min at room temperature, resulting in the formation of two liquid layers. The upper layer was transferred to a new glass vial using a glass Pasteur pipette. Lipids were extracted again from the lower layer by adding 300 µL MTBE, 100 µL MeOH and 100 µL of H2O, followed by shaking and centrifugation as in the first extraction. The upper layer from the second extraction was pooled with the upper layer from the first extraction and dried under nitrogen. Dried samples were reconstituted in 250 μL of 10 mM ammonium acetate in 50:50 MeOH:dichloromethane. A BCA assay was used to determine protein concentration in sample precipitate. Lipid levels were then normalized to protein content.

### Lipidomics – Mass spectrometry

Lipid and fatty acid species in the extracted cell samples were measured on a Sciex Lipidyzer mass spectrometry platform as previously described (29). The system consists of Shimadzu Nexera X2 LC-30AD pumps, a Shimadzu Nexera X2 SIL-30AC autosampler, and a Sciex QTRAP 5500 mass spectrometer equipped with SelexION for differential mobility spectrometry (DMS). 1-propanol was used as the chemical modifier for the DMS. Samples were introduced to the mass spectrometer by flow injection analysis at 8 µL/min. Each sample was injected twice, once with the DMS on (PC,PE,LPC,LPE and SM), and once with the DMS off (CE, CER, DAG, DCER, FFA, HCER, LCER, and TAG). Lipid molecular species were measured using multiple reaction monitoring (MRM) and positive/negative polarity switching. SM, DAG, CE, CER, DCER, HCER, DCER, and TAG were ionized in positive mode and LPE, LPC, PC, PE, and FFA in negative ionization mode, respectively. Data were acquired and processed using Analyst 1.6.3 and Lipidomics Workflow Manager 1.0.5.0. 440-665 lipids were measured across the study set of 6 samples. All the samples were prepared and analyzed in a single batch. For quality control, a pooled sample was run at the beginning and at the end of the batch, respectively. The median CV was 5.0%.

### iPSC-RPE and hfRPE culture

The generation and differentiation of normal and SFD iPSC-RPE lines in this manuscript have previously been described (15, 30). Informed consent was obtained from all subjects (University of Washington IRB-approved STUDY00010851). Fetal RPE were obtained from the Birth Defects Research Laboratory at the University of Washington (UW) under an approved protocol (UW5R24HD000836). All experiments were conducted according to the principles expressed in the Declaration of Helsinki.

Below we summarize how differentiated cells were cultured in this study. For the first week in culture, differentiated RPE cells were cultured in MEM-α medium (Invitrogen, 12561-072) supplemented with 5% (v/v) fetal bovine serum (FBS) (Atlanta Biologicals, S11550), 1x non-essential amino acid (NEAA) solution (Invitrogen, 11140-050), 1% (v/v) N1 supplement (Sigma-Aldrich, N6530), 50 U/mL penicillin/streptomycin (Invitrogen, 15070-063), 0.5 mg/L taurine (Sigma-Aldrich, T0625), 40 ng/L hydrocortisone (Sigma-Aldrich, H0396), 1.3 ng/L triiodo- thyronine at (Sigma-Aldrich, T5516), and 10 μM Y-27632 (Selleck chem, S1049). After the first week, FBS concentration was reduced to 1% FBS and Y-27632 was omitted for the rest of the culture period. Medium was changed once every 2-3 days.

A subset of iPSC-RPE and hfRPE cells were plated in sterile tissue culture-treated 96 well plates (Corning, 3603) at 50,000 cells/well (corresponding to data in **fig. 2, 3, 4f, 4h, 4i, 5a-c**). Cells were cultured for 4-6 weeks prior to experiments. Cells were supplied with 200 μL of medium per well. For experiments where cells were incubated in a ^13^C-labeled substrate, culture medium was serum-free but otherwise identical to conventional culture medium. For 96-well plates, medium volume was 200 μL/well.

**Figure 1.**
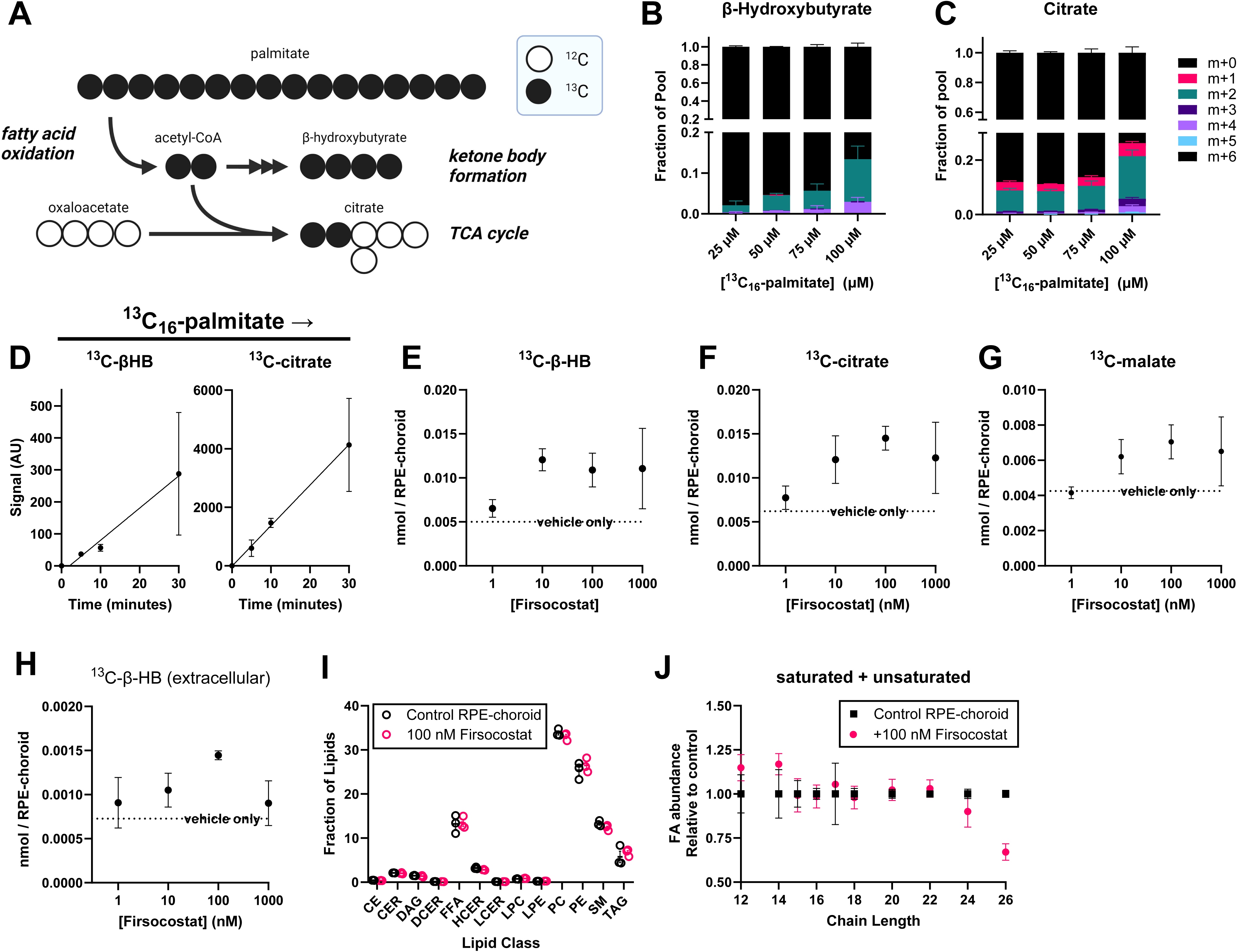
Firsocostat increases fatty acid oxidation in mouse RPE-choroid tissue. **(a)** The oxidation of a ^13^C16-palmitate molecule in mitochondria results in eight acetyl-CoA molecules, each with two ^13^C atoms (m+2 labeled). Each acetyl-CoA can enter the Krebs cycle to make m+2 citrate or can be used to synthesize ketone bodies. Because ketone bodies are synthesized from two acetyl-CoA molecules, they can be m+2 or m+4. **(b)** Increasing the extracellular concentration of ^13^C16-palmitate results in greater m+2 and m+4 labeling of β- hydroxybutyrate (β-HB) (n=3). **(c)** ^13^C accumulation on β-HB over 30 minutes is suppressed by the CPT inhibitor etomoxir (n=3). **(d)** ^13^C accumulates on β-HB and is exported to KRB buffer linearly with time (n=3). Application of Firsocostat to RPE-choroid for 1 hour increases the accumulation of ^13^C from palmitate on **(e)** intracellular β-HB, **(f)** intracellular citrate, **(g)** inctracellular malate, and **(h)** extracellular β-HB. The effect of Firsocostat is concentration-dependent, with a maximal effect at ∼100 nM (n=3). **(i)** 100 nM Firsocostat does not alter the distribution of lipid classes, though when fatty acid tails are summed across all classes, 100 nM Firsocostat appears to **(j)** decrease the proportion of very long chain fatty acid tails (n=3). CE: cholesterol esters, CER: ceramides, DAG: diacylglycerols, DCER: dihydroceramides, FFA: free fatty acids, HCER: hexosylceramides, LCER: lactosylceramides, LPC: lysophosphatidylcholines, PC: phosphotidylcholines, PE: phosphatidylethanolamines, SM: spingomyelins, TAG: triacylglycerols.

**Figure 2.**
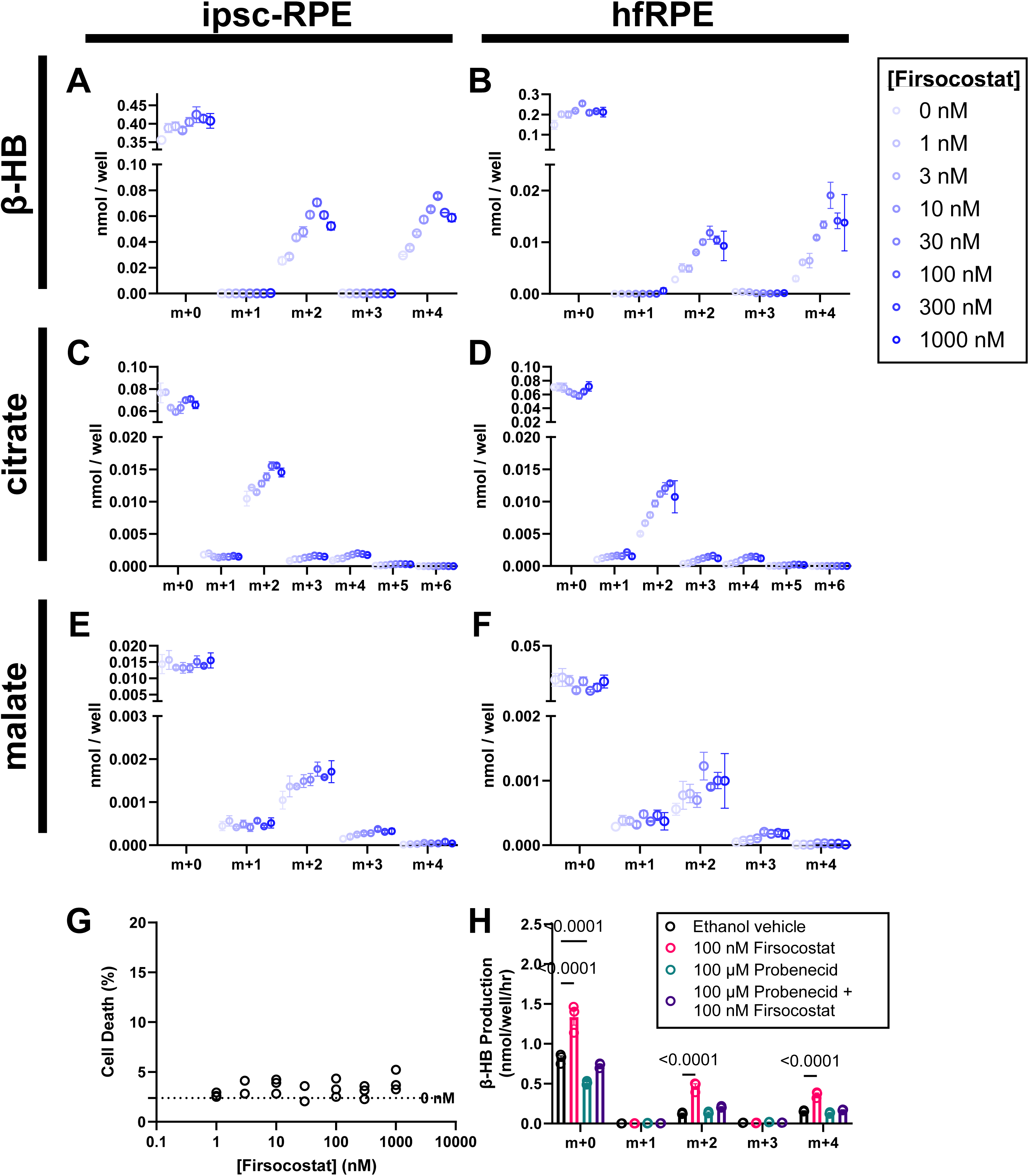
Firsocostat increases fatty acid oxidation in human RPE cells. iPSC-RPE cells **(a, c, e)** and human fetal RPE cells cultured on a 96-well plate **(b, d, f)** were supplied serum free medium supplemented with 50 µM ^13^C16-palmitate and 0, 1, 3, 10, 30, 100, 300 or 1000 nM Firsocostat (n=3). Six hours later we collected medium to determine ^13^C labeling on β-HB **(a, b)**, citrate **(c, d)**, malate **(e, f)**. Each isotopologue of a given metabolite is displayed separately. Darker blue circles correspond to higher concentrations of Firsocostat. **(g)** None of the concentrations of Firsocostat used significantly increased cell death, measured by LDH release into culture medium (n=3). **(h)** The Firsocostat-mediated increase in β-HB labeling was inhibited by probenecid, a non-selective inhibitor of organic anion transport proteins (n=4).

**Figure 3.**
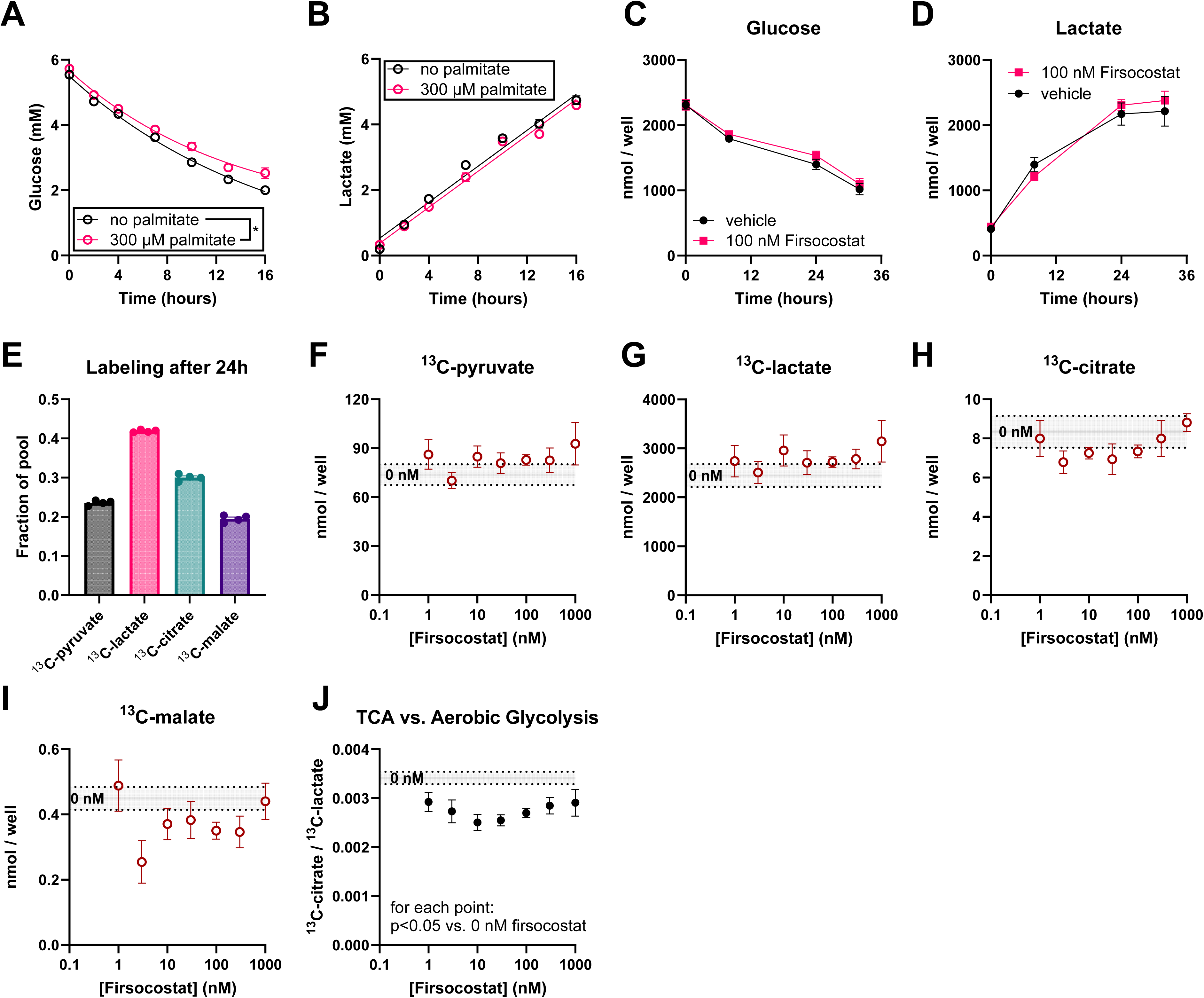
Firsocostat decreases glucose utilization by the TCA cycle. iPSC-RPE cells cultured on transwell filters were supplied with fresh Miller medium with or without 300 µM unlabeled palmitate. Medium was sampled 0, 2, 4, 7, 10, 13, and 16 hours after the medium change, then enzymatically assessed for (**a**) glucose and (**b**) lactate concentration. iPSC-RPE cells cultured on a 96-well plate were supplied with fresh Miller medium (containing 5 mM unlabeled glucose), with medium being sampled at 0, 8, 24, and 32 hours. **(c)** glucose and **(d)** lactate were assessed using enzymatic assays (n=6). There was no clear difference between glucose and lactate levels over time. To determine how glucose is being used, miller medium was supplemented with 5 mM ^13^C6-glucose and 0, 1, 3, 10, 30, 100, 300, 1000 Firsocostat (n=4). Medium was collected after 24 hours and we assessed ^13^C labeling of metabolites released by RPE. **(e)** In vehicle only samples, glycolytic and TCA cycle intermediates are well-labeled with ^13^C (maximum possible labeling is 50% because only 50% of glucose is ^13^C6). We determined the effect of Firsocostat concentration on all ^13^C labeled isotopologues of **(f)** pyruvate, **(g)** lactate, **(h)** citrate, **(i)** malate, and **(j)** the ratio of labeled citrate to labeled lactate. For all figures, the gray line indicates vehicle-treated controls, and dotted lines are the upper and lower bounds of the vehicle mean ± SEM.

**Figure 4.**
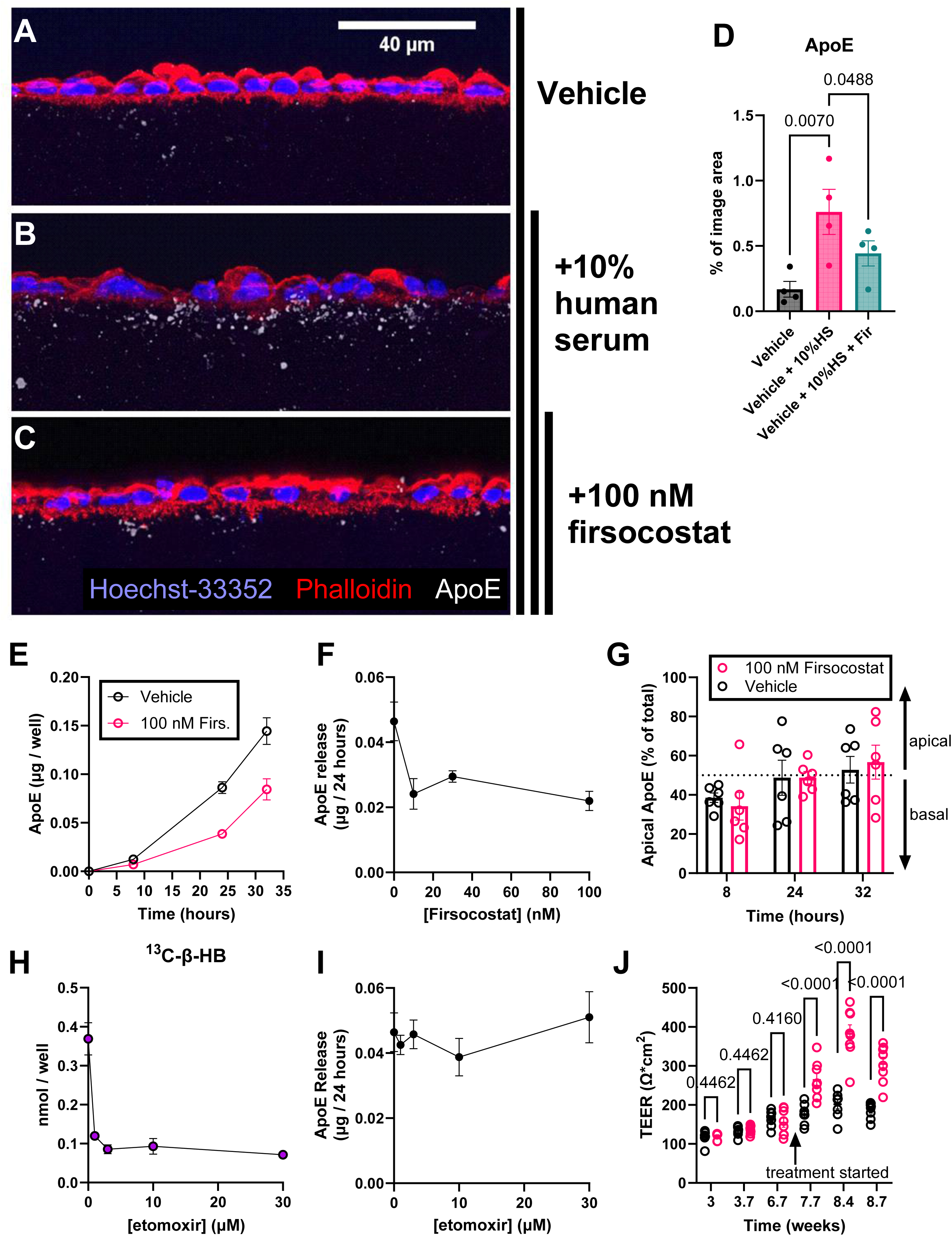
Firsocostat decreases ApoE accumulation. iPSC-RPE were grown on mixed cellulose ester membranes for 8 weeks, then the apical chamber was supplied with **(a)** medium containing vehicle, **(b)** vehicle and 10% human serum (v/v), or **(c)** 100 nM Firsocostat and 10% human serum. Firsocostat was provided to both apical and basal chambers. Cells were fixed in 4% paraformaldehyde and labeled to visualize nuclei (Hoechst-33352, blue), F-actin (phalloidin, red), and ApoE (white). Human serum causes the formation of ApoE puncta in the membrane below RPE cells. **(d)** The area of the membrane occupied by ApoE puncta is decreased with Firsocostat (n=4). ApoE is normally released by RPE cells over time, and **(e)** 100 nM Firsocostat decreases the rate of ApoE release (sum of ApoE in apical and basal chambers, n=6). **(f)** Over a 24-hour period, 10 nM, 30 nM, and 100 nM Firsocostat all decrease ApoE release to a similar extent (n=4). **(g)** ApoE release is not polarized to the apical or basal side, and Firsocostat does not change the polarity of ApoE release (n=6). **(h)** etomoxir decreases the ability of ^13^C16- palmitate to form ^13^C-βHB, yet **(i)** the etomoxir-mediated decrease in fatty acid oxidation does not alter ApoE release over 24 hours (n=4). **(j)** When provided to mature RPE (after 6.7 weeks in culture), Firsocostat increased TEER (n=8).

RPE cells in other experiments (**Fig. 4a-e, 4g, 4j, 5d-f**) were cultured in 24-well plates (Falcon, 353047) on mixed cellulose ester Millicell inserts (Millipore, PIHA01250). Cells were plated at 100,000 cells/insert and cultured 6-8 weeks before experimental manipulations. We supplied cells with 400 μL of culture medium to the apical chamber and 600 μL to the basal chamber of each well. RPE cells used for brightfield imaging (**Fig. 5g**) were cultured on clear polyester culture inserts (Corning, 3450) on 24-well plates . These cells were treated identically to RPE cultured on mixed cellulose ester inserts.

**Figure 5.**
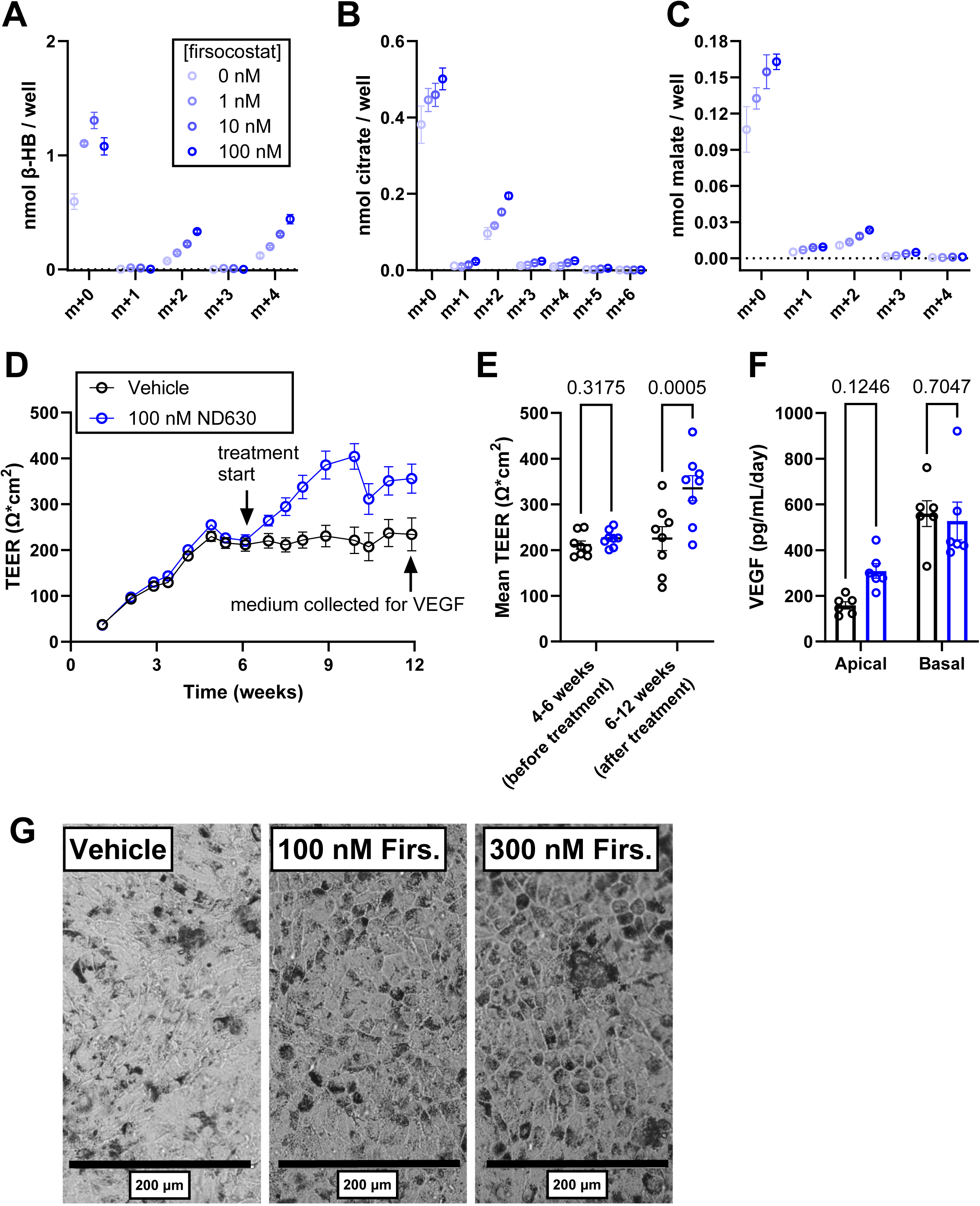
Firsocostat increases fatty acid oxidation and enhances barrier function in iPSC-RPE from donors with Sorsby’s fundus dystrophy (SFD). iPSC-RPE cells from donors with SFD were cultured on a 96 well plate. At 6 weeks in culture, cells were provided with 50 µM ^13^C16-palmitate and 0 nM, 1 nM, 10 nM, or 100 nM for 8 hours (n=4). Following the incubation, culture medium was collected and we determined ^13^C labeling on **(a)** βHB, **(b)** citrate, and **(c)** malate. Each of these intermediates is downstream of fatty acid oxidation. RPE was also cultured on polyester filters, either with vehicle, 100 nM Firsocostat, or 300 nM Firsocostat. **(d)** As with iPSC-RPE from normal donors, 100 nM Firsocostat increases TEER (n=8). **(e)** Release of VEGF by vehicle- and Firsocostat-treated SFD-iPSC-RPE cells is polarized such that more is release to the basal side, but apical VEGF release is selectively increased in cells treated with 100 nM Firsocostat (n=6). **(f)** Firsocostat appears to increase the proportion of SFD-iPSC-RPE that are polygonal in shape.

For immunostaining experiments, cells were treated with vehicle or 10% (v/v) normal human serum (Millipore, S1-100ML, lot# 2330486) in serum free media, supplied to the apical chamber at 8 week for 48 hours. When Firsocostat (Caymen, 23961) was supplied to cells, it was made as a 100x solution in ethanol and diluted in medium. When probenecid (Caymen, 14981) was supplied to cells, it was diluted from a 100x stock in DMSO, and DMSO was also used in the control group.

### Trans-epithelial electrical resistance (TEER)

We measured TEER in of RPE plated on transwell filter membranes using an Epithelial Volt-Ohm Meter (Millipore, #MERS00002) and ERS Probes (Millipore, #MERSSTX01). TEER was measured in triplicate for each well and averaged (*TEERwell*). A cell-free TEER which only includes the filter membrane (*TEERblank*) is also determined and subtracted from *TEERwell*. The resulting TEER (in Ω) is multiplied by the culture surface area *Areamembrane* to obtain final resistance values. For mixed cellulose ester filters, *Areamembrane* = 0.6 cm^2^.

### Enzyme-linked immunosorbent assays (ELISAs)

For ELISA experiments, cells were treated with vehicle (0.1% ethanol) or indicated concentrations of firsocostat at 6 weeks of age with medium being changed as indicated above.

After two weeks of the treatment, the 50 μL of media was collected at 0, 8, 24 and 36 hours (figure 4E), and 24 hours post treatment (figure 4F) for ApoE ELISA. Similarly, the 50 μL of medium was collected after 24 hours post treatment for VEGF ELISA.

We used ELISAs to quantify human ApoA1 (R&D biosystems, DY3664), ApoB (R&D biosystems, DAPB00), ApoE (Abcam, ab108813), and VEGF (R&D biosystems, DY293B) content in cell culture supernatant. For each assay we followed the manufacturer’s instructions, and all quantified data were in the linear range of the standard curve. Absorbance was measured on a BioTek Synergy 4 plate reader.

### Glucose and lactate assays

Glucose and lactate levels in cell culture supernatant were measured using previously described enzymatic assays (31) adapted for use in a 96 well plate. A detailed description of the adapted glucose assay protocol can be found at https://www.protocols.io/view/glucose-concentration-assay-hexokinase-g6pdh-metho-dm6gpj5jdgzp/v1. A detailed description of the adapted lactate assay can be found at https://www.protocols.io/view/lactate-concentration-assay-ldh-method-6qpvr4733gmk/v1. For each assay, enzymatic consumption of glucose or lactate is linked to NAD(P)H production. We determined the correspondence between glucose or lactate content and NAD(P)H by measuring absorbance at 340 nm by using freshly prepared standard curves. Absorbance was measured using a BioTek Synergy 4 plate reader (Agilent).

### Lactate dehydrogenase assays

We quantified LDH levels using a CyQuant^TM^ Cytotoxicity Assay kit (ThermoFisher C20300) and followed the manufacturers protocol. Positive LDH controls were fully lysed in 10% Triton-X100, negative controls were cell-free culture medium. We confirmed that all absorbance measurements were in the linear range of the reaction using a BioTek Synergy 4 plate reader.

### Sectioning and Immunofluorescent Labeling

For immunohistochemical analysis of protein localization, cells were fixed using 4% paraformaldehyde in 1x phosphate buffered saline (PBS, pH 7.4) for 24 hours. Filters were then washed in PBS and quartered. Filter quarters were embedded in OCT then sectioned at -20°C at a 20 µm thickness. Sections were stored at -80°C.

Filter sections were heated for 30 minutes to adhere filters to slides, and an oil pen was used to form a hydrophobic barrier on slides. Slides were washed in 1x PBS, pH7.4, non-specific antigens blocked using 2% donkey serum and 0.3% Triton X-100 in PBS, incubated in primary cocktail made of goat α-ApoE (EMD-Millipore, AB947, 1:500), Hoechst-33328 (ThermoFischer, H3570, 5 µg/mL), and phalloidin-568 (molecular probes, A12380, 2U/mL) overnight at 4°C, washed three times, and labeled with secondary antibody (AlexaFluor633-conjugated donkey α-goat, Invitrogen, A21082, 1:1000) for 2 hours at room temperature. Fluorescent sections were washed twice more, mounted in fluoromount G, and sealed with clear nail polish. Slides were imaged on a Leica SP8 confocal microscope. Each image covered a 144.87 µm (x) by 144.87 µm (y) area (at 1024x1024 pixels). For each imaging area, eleven z-stack images were taken covering a 10 µm thickness. 9-10 replicate images were taken from different regions of each RPE mixed cellulose ester membrane.

### Image Analysis

ApoE puncta are quantified in **Fig. 4d**. Image processing and puncta quantification used ImageJ (v1.53t). Image stacks were made into a maximum intensity projection prior to downstream analysis. The ApoE channel of this projection was processed with a gaussian blur function (sigma=1). Background was subtracted using a sliding paraboloid function, then an intensity threshold was applied (30–255). The number, size, and coverage of the ApoE channel were determined using the “analyze particles” tool. A flow chart of the analysis performed on a sample image is available as **supplemental Fig. 1**. Images displayed in **Fig. 4a-c** are background subtracted. Particle analysis results were averaged, such that a single point represents the average of 9-10 images made across that membrane. Different replicates represent different membranes.

### Statistical analysis

All displayed include mean ± standard error and may additionally show individual experimental replicates. Statistical analyses were performed using Prism v10.0 (GraphPad Software).

Significance thresholds were set to α<0.05. When below this threshold, statistical values are noted within the figure. Statistical tests are as follows: <u>Student’s t-test</u>: Fig. 1c; <u>1-way ANOVA</u>: Fig. 1E-H, 2G, 3D-H, 4D; <u>2-way ANOVA</u>: Fig. 1I, 1J, 2H, 4G, 4J, 5E, 5F. All post-hoc hypotheses were tested using the Two-stage linear step-up procedure of Benjamini, Krieger and Yekutieli.

## Results

### Firsocostat increases fatty acid oxidation by mouse RPE-choroid-sclera

Many experiments in this manuscript rely on assessing fatty acid oxidation (FAO) of ^13^C16- palmitate (all carbons of the palmitate molecule carry the ^13^C isotope). Acetyl-CoA from oxidized ^13^C16-palmitate is two atomic mass units heavier (m+2) than normal acetyl-CoA, so when ^13^C16-palmitate-derived acetyl-CoA is incorporated into citrate (in the Krebs cycle) or β- hydroxybutyrate (β-HB; a ketone body), each should be m+2 or m+4 labeled due to incorporation of one or two ^13^C-labeled acetyl-CoA molecules (**Fig. 1a**).

We determined whether mouse RPE-choroid performs sufficient fatty acid oxidation (FAO) to detect labeling of FAO products. We provided RPE-choroid with 25-100 µM ^13^C16-palmitate in KRB + 5 mM unlabeled glucose for 30 minutes. There is a concentration-dependent increase in labeling of tissue β-HB (**Fig. 1b**), which was predominantly m+0, m+2, or m+4 labeled – supporting the hypothesis that the ^13^C is derived from β-oxidation of ^13^C16-palmitate to ^13^C- acetyl-CoA. Oxidation of long-chain fatty acids such as palmitate requires carnitine acyltransferases, which are inhibited by etomoxir (32). Etomoxir reduces the accumulation of ^13^C from 50 µM ^13^C16-palmitate in RPE-choroid tissue over 30 minutes (**Fig. 1c**). The generation of ^13^C-β-HB and export into medium is linear over time (**Fig. 1d**). These data show that we can assess oxidation of long-chain free fatty acids by mouse RPE-choroid.

Malonyl-CoA produced by ACCs inhibits carnitine acylation to long-chain free fatty acids and thus their import into mitochondria for β-oxidation (21). ACCs are inhibited by the small molecule Firsocostat. By blocking production of malonyl-CoA Firsocostat dis-inhibits FAO in hepatocytes (23). We determined whether Firsocostat produces the same effect in mouse RPE-choroid-sclera. 100 nM Firsocostat increases the generation and export of ^13^C- β-HB from 50 µM ^13^C16- palmitate (**Fig. 1d**). We incubated freshly dissected tissue in 50 µM ^13^C16-palmitate, 5 mM unlabeled glucose and 0, 1, 10, 100, or 1000 nM Firsocostat for 60 minutes. Firsocostat increased ^13^C labeling on intracellular β-HB (**Fig. 1e**), citrate (**Fig. 1f**), and malate (**Fig. 1g**). ^13^C labeled β-HB was also found in culture medium (**Fig. 1h**), suggesting it is produced and exported. Fractional labeling of β-HB in tissue and culture medium are similar but labeling is more dilute in medium, due to the presence of unlabeled β-HB stores that are produced and exported prior to the tissue oxidizing ^13^C16-palmitate. Notably, retinas are also able to oxidize ^13^C16-palmitate, but unlike RPE-choroid their ability to do so is not enhanced by Firsocostat (**supplemental Fig. 2**)

We determined whether lipid profiles in RPE-choroid are also affected by Firsocostat. We exposed freshly dissected mouse RPE-choroid-sclera to 50 µM unlabeled palmitate, 5 mM glucose, and either vehicle or 100 nM Firsocostat in KRB for 6 hours. Tissue was flash-frozen and lipid profiles assessed on the Lipidyzer mass spectrometry platform. Firsocostat did not appear to alter lipid class (**Fig. 1i**) but does affect the distribution of fatty acid tails on lipids, with a decrease in very long-chain fatty acid tails (C26) and an increase in species carrying short chain fatty acid tails that is not statistically significant (**Fig. 1j**). These data were initially unexpected because an increase in mitochondrial fatty acid oxidation should selectively affect fatty acids with chain lengths ≤20 carbons. However, the time required for changes in fatty acid oxidation to affect total intracellular free- and lipid-bound fatty acid composition (33) may be sufficiently slow that tissues require a longer exposure to the drug to observe an effect of Firsocostat on lipid profiles. Additionally, malonyl-CoA is able to inhibit peroxisomal carnitine acyltransferases (34), so Firsocostat may also impact peroxisomal lipid metabolism.

### ACC inhibition stimulates fatty acid oxidation by cultured human RPE cells

RPE-choroid-sclera tissue contains RPE cells but also a constellation of other cell types (35). RPE cells are considered a top candidate among the cell types that could produce drusen. In culture, RPE cells are sufficient to produce drusen-like deposits that may be quantified to assess the level of AMD-like pathology (36–38). We determined whether Firsocostat also increases FAO in cultures of pure RPE cells. We treated both iPSC-RPE cells (**Fig. 2a, 2c, 2e**) derived from normal subjects and human fetal RPE (hfRPE; **Fig. 2b, 2d, 2f**) with 1-1000 nM Firsocostat in RPE culture medium containing 50 µM ^13^C16-palmitate. Over 6 hours, Firsocostat increases accumulation of ^13^C on β-HB (**Fig. 2a-b**), citrate (**Fig. 2c-d**), and malate (**Fig. 2e-f**) that are released in culture medium. Concentrations exceeding 100 nM had either no effect or caused a decline in ^13^C labeling. The decline in ^13^C labeling does not appear to result from Firsocostat toxicity, as a 24 hour exposure to Firsocostat does not appreciably increase cell death (**Fig. 2g**). Firsocostat is designed to act upon tissues that express organic anion transport proteins (OATPs) (39). These transporters are strongly expressed on hepatocytes and are inhibited by probenecid. 100 µM probenecid abolishes the Firsocostat-mediated increase in flux of ^13^C from palmitate to β-HB in iPSC-RPE (**Fig. 2h**), suggesting that RPE cells also express a member of the OATP transporter family, which is used to import the drug.

### Firsocostat-dependent effects on glucose metabolism

When Firsocostat increases FAO it provides more energy from fatty acids. If cellular energy demand is unchanged by Firsocostat, there may be a compensatory decrease in activity of other energetic pathways. In other tissues, inhibition of glucose consumption by fatty acid metabolism is referred to as the Randle cycle (40). We determined whether additional fatty acid content in medium or Firsocostat alter glycolysis or glucose-dependent Krebs cycle activity in iPSC-RPE cells. In our first experiment, we added 300 µM palmitate to medium and determined (**Fig. 3a**) glucose and (**Fig. 3b**) lactate concentrations 0, 2, 4, 7, 10, 13, and 16 hours following the medium change. The best fit parameters for the glucose uptake data were statistically different with vs. without palmitate, suggesting that glucose uptake is slightly impaired in the presence of palmitate.

We also monitored changes in medium glucose and lactate levels over a 32 hour period when iPSC-RPE were treated with 50 µM palmitate in the presence of vehicle or 100 nM Firsocostat. Glucose consumption (**Fig. 3c**) and lactate production (**Fig. 3d**) are unaltered by Firsocostat. We also probed medium from RPE cells supplied with 50% labeled ^13^C6-glucose for 24 hours. Glucose labels lactate, pyruvate, citrate, and malate in culture medium well (**Fig. 3e**), suggesting that changes in the utilization of glucose to make these metabolites should be clear. On average groups treated with Firsocostat made more ^13^C-labeled pyruvate (**Fig. 3f**) and lactate (**Fig. 3g**) while making less citrate (**Fig. 3h**) and malate (**Fig. 3i**). The variation between samples is greater than the effect size, so none of these effects are statistically significant. We

normalized levels of ^13^C-labeled citrate to levels of ^13^C-labeled lactate from the same sample, thus quantifying the ratio of carbons from glucose taking the Krebs cycle route vs aerobic glycolysis. This normalization reduces sample-to sample variation, and revealed a small, statistically significant influence of Firsocostat on glucose utilization (**Fig. 3j**). Together the data show that Firsocostat does not alter the rate of glucose consumption, but it slightly diminishes glucose input into the Krebs cycle relative to lactate formation. This implies that the Krebs cycle may compensate for increased fatty acid oxidation by slightly diminishing Krebs cycle input form glucose.

### Firsocostat remodels lipoprotein transport

Since lipoproteins can contribute to drusen we determined whether Firsocostat can influence accumulation of lipoprotein particles in Bruch’s membrane. Lipid and apolipoprotein E (ApoE) deposition within filter substrates on which iPSC-RPE cells are grown approximate the phenotype of AMD-like pathologies in culture (36, 37). Human serum stimulates the accumulation of ApoE-containing particles in these filters (36).

Firsocostat decreases accumulation of basal ApoE deposits. We supplemented culture medium in the apical chamber with 10% human serum for 48 hours, washed the RPE cells, and then fixed them in 4% paraformaldehyde. We immunolabeled cryosections of these filters to detect ApoE within the filter. ApoE particles are visible in all treatment groups (**Fig. 4a-c**). 10% human serum increases the cross-sectional area of the filter occupied by ApoE deposits (**Fig. 4a-b, d**). 100 nM Firsocostat blunts the effect of 10% human serum so that ApoE deposits occupy a smaller area of the filter (**Fig. 4b-d**). We confirmed this finding with iPSC-RPE cultures treated with or without 100 nM Firsocostat by using an ELISA assay to quantify release of ApoE into the culture medium. Firsocostat significantly diminishes the release of ApoE (**Fig. 4e**). These experiments show that Firsocostat can influence the rate of free fatty acid oxidation (FAO), lipid metabolism, and lipoprotein release. This effect is consistent across a range of Firsocostat concentrations (**Fig. 4f**) and is not polarized to either the apical or basolateral sides (**Fig. 4g**). Our findings imply that increasing FAO deprives RPE of sufficient lipid to support the assembly and export of lipoprotein particles.

Since increased FAO suppresses lipoprotein release, we asked whether inhibiting FAO stimulates lipoprotein release. We treated RPE cells with 50 µM ^13^C16-palmitate-BSA and increasing concentrations (0, 1, 3, 10, 30 µM) of the carnitine acyltransferase inhibitor etomoxir. Etomoxir inhibits oxidation of palmitate in iPSC-RPE mitochondria (**Fig. 4h**) but it does not alter ApoE release (**Fig. 4i**). This suggests that rates of FAO alone do not regulate ApoE release, or that RPE normally releases the maximal amount of ApoE it is able to. Long-term Firsocostat treatment also improves tight junction formation, as RPE cell transepithelial electrical resistance (TEER) increases significantly after treatment with Firsocostat (**Fig. 4j**).

Lipoproteins may be assembled within RPE cells from endogenous sources of fatty acids, cholesterol and ApoE and then exported. Other types of lipoproteins may pass through the RPE from the circulation on the basolateral side to the retina on the apical side and be remodeled along the way. Firsocostat-treatment of RPE cells could affect the remodeling of those lipoproteins. ApoA1 associates with high-density lipoproteins (41), while ApoB100 associates with low density lipoproteins (42). We evaluated effects of Firsocostat on lipoprotein transport across the RPE by providing iPSC-RPE on transwell filters with 10% human serum in the basal chamber to mimic the supply of lipoprotein particles that would come from circulation. If Firsocostat remodels lipoproteins, the rate or extent to which they reach the apical chamber would be altered. We found that Firsocostat diminishes accumulation of ApoA1 in the apical chamber (**supplemental Fig. 3a**), whereas it does not influence ApoB100 transport (**supplemental Fig. 3b).** Firsocostat does not alter the overall rate of cholesterol transport (both esterified and unesterified; **supplemental Fig. 3c**). These data suggest that Firsocostat does not affect bulk lipoprotein transport from the choroid through the RPE to the retina, but it may affect the composition of lipoproteins that reach the retina.

### Firsocostat improves cellular physiology of iPSC-RPE cells derived from a patient with Sorsby’s fundus dystrophy

Sorsby’s fundus dystrophy (SFD) is a severe inherited retinal disorder with clinical and histological characteristics that overlap with AMD. SFD patients have a thicker Bruch’s membrane, which accumulates large, lipid-rich sub-RPE deposits (15, 43). SFD is caused by mutations in an extracellular matrix metalloprotease inhibitor TIMP3 (44), but the relationship between lipid pathology and the TIMP3 mutations is unclear.

In iPSC-RPE generated from a SFD patient, Firsocostat stimulates oxidation of 50 µM ^13^C16- palmitate-BSA to make ^13^C-citrate (**Fig. 5a**), ^13^C-BHB (**Fig. 5b)**, and ^13^C-malate (**Fig. 5c**) that accumulate in culture medium after 8 hours. As it does with iPSC RPE from normal donors, addition of Firsocostat to SFD-iPSC-RPE increases TEER for at least 6 weeks (**Fig. 5d**). We quantified the steady-state TEER for each well before and after application of 100 nM Firsocostat. Fig. 5e shows that 100 nM Firsocostat increases TEER. After 12 weeks we checked cell polarity by quantifying VEGF released from RPE to the apical and basal chambers. SFD RPE, like normal RPE, secretes more VEGF into the basal chamber (15), (**Fig. 5f**).

Notably, 100 nM Firsocostat increases apical VEGF release. An increase in VEGF release could be detrimental if it encourages aberrant inner retinal blood vessel growth, but VEGF signaling is pleiotropic and in addition to encouraging vessel growth can also sustain Müller glia and inner retinal neurons without impacting blood vessel integrity (45). Finally, Firsocostat improves iPSC-SFD-RPE cellular morphology. RPE cells normally are hexagonal, so this is a desirable trait in culture. A loss in hexagonal morphology can be a sign that RPE have undergone an epithelial-to-mesenchymal transition (46). 100 or 300 nM Firsocostat substantially improves the hexagonal character of iPSC-SFD-RPE (**Fig. 5g**). Together these data imply that Firsocostat not only stimulates fatty acid oxidation, but also enhances the overall morphology and physiology of RPE cells.

## Discussion

There have yet to be any FDA-approved treatments for geographic atrophy that directly affect lipid metabolism, yet the need for drugs that target lipid and lipoprotein accumulation has been acknowledged by leaders in the field (16, 17). Our investigation of ACC inhibition begins to address that need.

Our studies show that Firsocostat accelerates fatty acid oxidation (FAO) in mouse RPE-choroid tissue and in cultured human RPE cells. Stimulating FAO in these cells influences glucose, lipid, and lipoprotein metabolism. Firsocostat slows the release of ApoE from RPE cells and it limits deposition of ApoE-containing particles within the filter substrate on which the RPE cells grow. We hypothesize that Firsocostat limits ApoE release because there is less intracellular lipid to release. If this hypothesis is correct then treatment of an AMD patient with Firsocostat could slow accumulation of lipoproteins within drusen. This model is depicted in **Fig. 6**. If Firsocostat can slow accumulation of drusen or thickening of Bruch’s membrane in patients, it may be possible to use Firsocostat treatment to determine whether these phenotypes contribute directly to photoreceptor degeneration and potentially to use Firsocostat or other ACC inhibitors to slow the progression of AMD.

**Figure 6.**
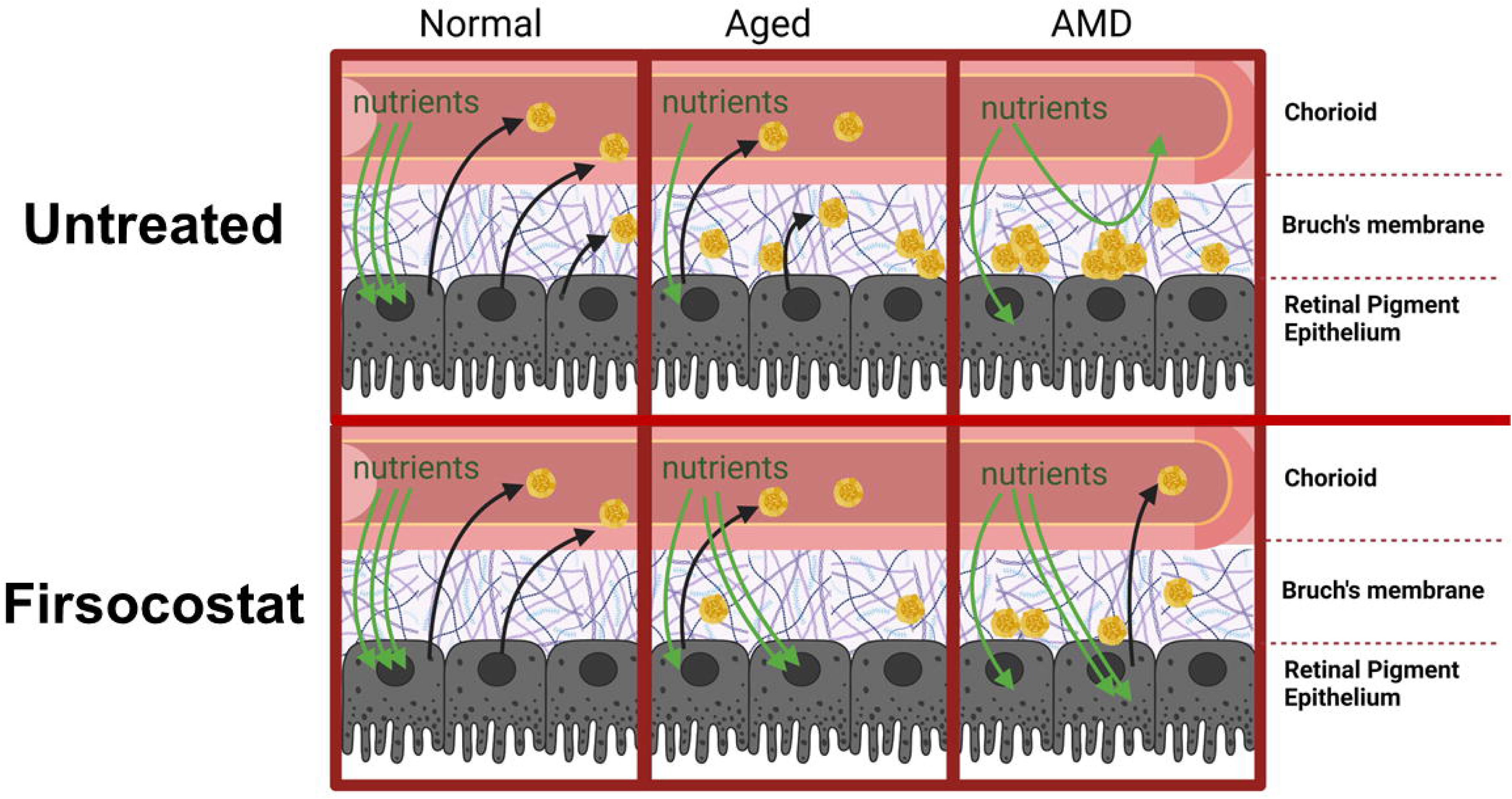
Diagram illustrating the hypothetical role Firsocostat may play in lessening AMD pathology.

Complement factors accumulate in drusen and activation of the complement system contributes to AMD progression. The only FDA-approved treatments for geographic atrophy (which occurs in advanced dry AMD) are inhibitors of the complement proteins C5 and C3 (47, 48). Notably the complement system and lipoproteins interact. Both complement factors and apolipoproteins accumulate in druse. We propose that Firsocostat treatment can diminish drusen by limiting release of lipoproteins from RPE cells. However, it will be important to determine if diminished levels of ApoE containing lipoproteins could have undesirable effects. For example, ApoE can inhibit the complement cascade directly by binding activated C1Q (49), so it will be important to determine if limiting the release of ApoE from RPE cells allows more activation of complement pathways (50).

Our studies so far have investigated effects of an ACC inhibitor in cultured cells and in isolated mouse RPE/choroid tissue. They are only a first step toward targeting metabolism to improve AMD outcomes. The next steps will be to determine (a) whether circulating Firsocostat or other ACC inhibitors enter RPE cells *in vivo*, (b) whether Firsocostat alters not only lipoprotein levels but also composition, (c) whether Firsocostat diminishes accumulation of sub-RPE lipid deposits in animal models of AMD, and (d) whether RPE lipid metabolism compensates for chronic administration of ACC inhibition. SREBP1c is activated by genetic ablation of ACC1 and ACC2 and can normalize or increase systemic lipid levels (51), so an important goal will be to determine how systemic lipid levels affect RPE lipoprotein accumulation.

## Acknowledgements

The authors would like to thank Susan Brockerhoff and Ian Sweet for helpful discussions, and Marcos Nazario for his role in maintaining iPSC-RPE cells. The Birth Defects Laboratory is supported by NIH award number 5R24HD000836 from the Eunice Kennedy Shriver National Institute of Child Health and Human Development. Patient-derived iPSC lines were generated by Chris Cavanaugh, Jennifer Hesson, and Carol Ware in the Ellison ISCRM stem cell core facility. Figure 1A and 6 were created with Biorender.com. Lipidomics were performed by the Northwest Metabolomics Research Center at the University of Washington, Seattle. The mass spectrometry system used to acquire quantitative lipidomics data was supported by NIH 1S10OD021562-01.

