## Supplementary figures and images for "Acetyl-CoA carboxylase Inhibition increases RPE cell fatty acid oxidation and limits apolipoprotein efflux"

### Supplemental Figure 1

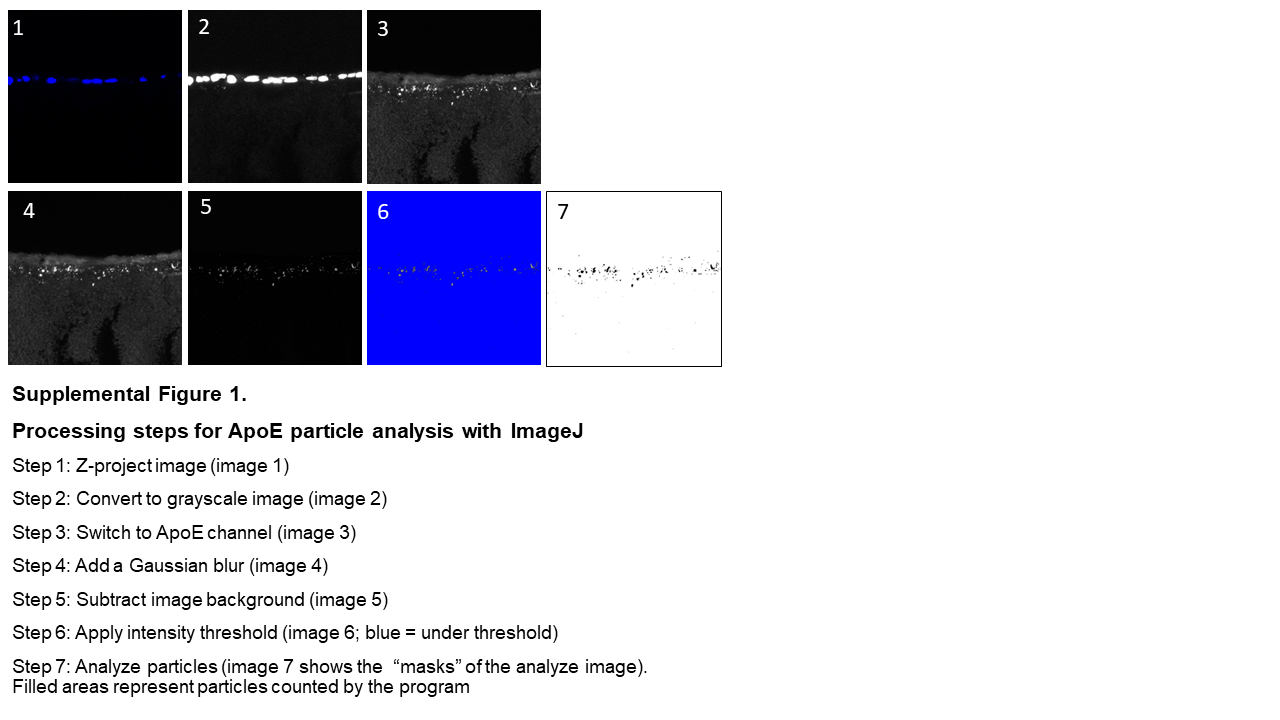

### Supplemental Figure 2

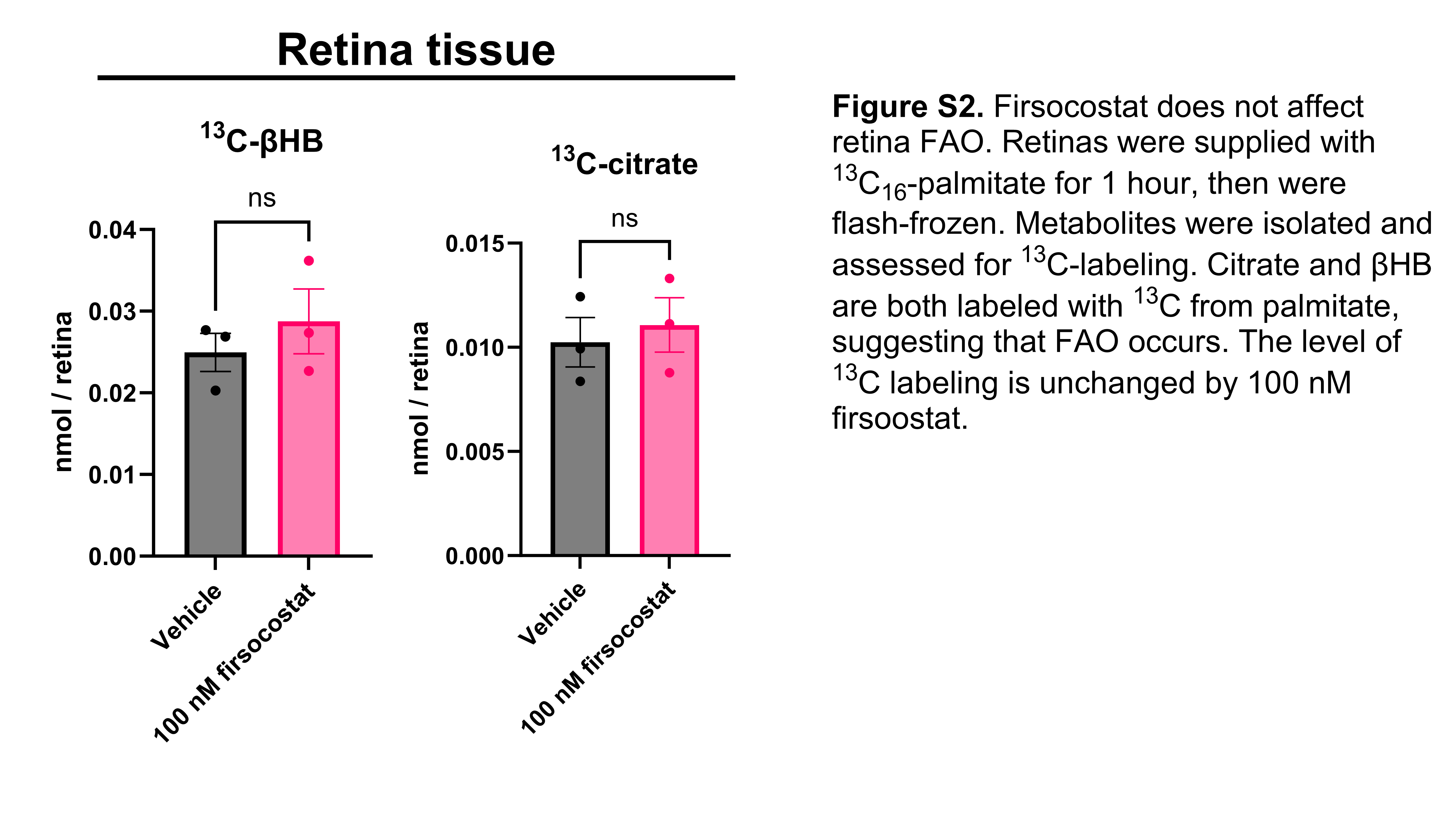

### Supplemental Figure 3

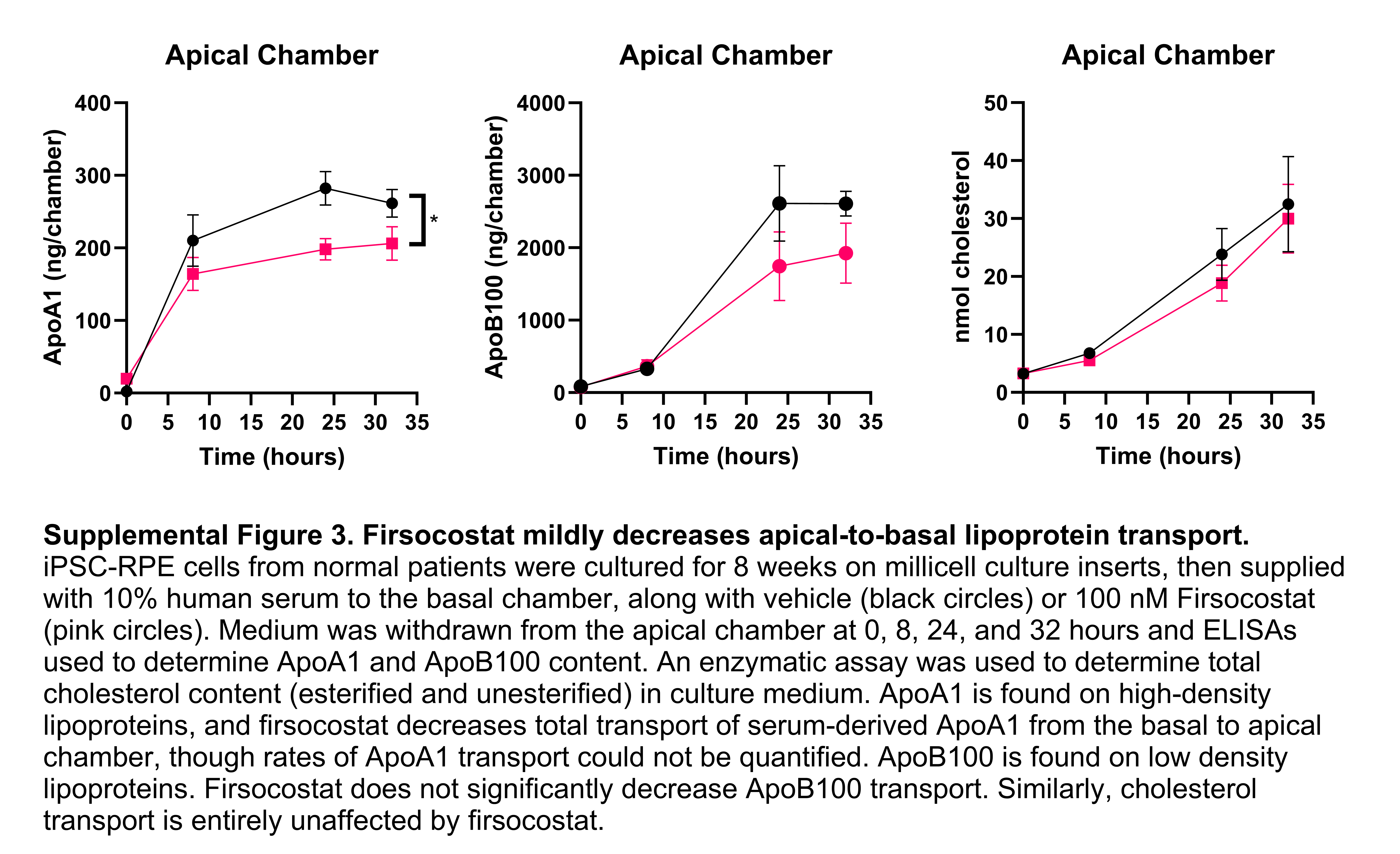
